# A comparison of two culture and three sequencing methods for detection of the urobiome

**DOI:** 10.64898/2026.10.28.762191

**Authors:** Mark Khemmani, Melline Fontes Noronha, Thythy T. Pham, Ben Ferneyhough, Matthew J. Roddis, Nick J. Parkinson, Alan J Wolfe

**Affiliations:** Department of Microbiology and Immunology, Stritch School of Medicine, Loyola University Chicago, Maywood, IL, USA; Division of Urogynecology & Reconstructive Pelvic Surgery, Department of Obstetrics & Gynecology, Stritch School of Medicine, Loyola University Chicago, Maywood, IL; Metagenomic Testing Company Ltd, Abingdon, Oxfordshire OX14 4SA, United Kingdom

**Keywords:** 16S rRNA gene sequencing, bacterial detection, expanded quantitative urine culture, microbiome, shotgun sequencing, standard urine culture, urinary tract, urobiome

## Abstract

Standard urine culture (SUC), the clinical reference method, is optimized for fast-growing aerobes and misses much of the urinary microbiota. Enhanced culture and sequencing methods have rarely been compared on the same specimens. We compared five methods (SUC, expanded quantitative urine culture [EQUC], short-read V4 16S rRNA gene sequencing, long-read near full-length 16S rRNA gene sequencing, and PCR-free long-read shotgun metagenomic sequencing) on parallel aliquots of urine from pregnant participants screened for asymptomatic bacteriuria. Of 98 cultured samples, 98 (100.0%) were EQUC-positive while 85 (86.7%) SUC-positive. EQUC detected 102 species, but SUC only detected 41. Short-read 16S rRNA gene sequencing, long-read 16S rRNA gene sequencing, and shotgun sequencing were positive for 188/188, 186/188 and 139/139 samples, detecting 195, 69 and 528 species, respectively. In the 79 samples positive by EQUC and all three sequencing methods, sequencing profiles resembled each other more than culture profiles, and SUC captured only a subset of taxa detected by every other method. Shotgun sequencing gave the broadest taxonomic coverage and was the only sequencing method to identify *Escherichia coli* to the species level. With EQUC, it outperformed both 16S sequencing methods in detecting *Gardnerella vaginalis* and distinguishing *Lactobacillus* species, and it resolved six *Gardnerella* species, three detected by no other method. Long-read 16S rRNA gene sequencing gave the most confident amplicon-based species assignments, whereas culture better recovered *Corynebacterium* and *Staphylococcus* species. No single method was best across all endpoints, but long-read shotgun metagenomics provided the broadest species-level view. Method choice should match the research or clinical question; outcome-based studies are needed to establish clinical utility.

**IMPORTANCE:** The microbes reported in a urine sample depend heavily on how that sample is analyzed. Standard urine culture, the test most clinical laboratories rely on, detected only a fraction of the bacteria found by enhanced culture or by DNA sequencing of the same specimens. Sequencing methods also differed from each other: short-read 16S rRNA gene sequencing captured broad diversity but often could not name species, whereas long-read methods distinguished clinically and ecologically important species. PCR-free long-read shotgun metagenomics detected the widest range of taxa and resolved species, such as *Escherichia coli* and multiple *Gardnerella* species, that amplicon methods could not. Because each method has distinct strengths and blind spots, detection methods should be chosen according to the question being asked, and sequencing-based diagnostics must be validated against patient outcomes before they can replace culture.

## INTRODUCTION

Urinary tract infection (UTI) is among the most common bacterial infections encountered in clinical practice, affecting a large proportion of adult females in their lifetime and imposing a substantial and rising global burden of morbidity, healthcare cost, and antibiotic use (Flores-Mireles et al. 2015). Accurate identification and quantification of microbes present in urine is fundamental to diagnosing UTI and other lower urinary disorders (*e.g.*, urgency urinary incontinence) and conditions, (*e.g.*, asymptomatic bacteriuria in pregnant females). Accurate detection also is critical for understanding the underlying microbiology. Yet the laboratory methods for generating that information vary widely in sensitivity, resolution and interpretability (Brubaker et al. 2023).

For more than half a century, the diagnostic mainstay has been standard urine culture (SUC), where mid-stream voided urine is plated under aerobic conditions and a uropathogen is reported if it exceeds a threshold, conventionally 10^5^ colony-forming units per milliliter (Kass 1957; Nicolle et al. 2019; Public 2019). While both rapid and inexpensive, this approach is optimized to detect fast-growing facultative anaerobic microbes (*e.g.*, *E. coli*) and systematically overlooks fastidious, slow-growing, and anaerobic bacteria, as well as clinically relevant microbes present below the reporting threshold (Moreland et al. 2024).

A substantial fraction of patients with typical UTI symptoms returns a “no growth” or negative standard culture; however, the criteria underpinning SUC have been increasingly characterized as a “tarnished gold standard” that no longer reflects contemporary understanding of the urinary tract (Price et al. 2017; Brubaker et al. 2023). Enhanced culture and molecular re-examination of culture-negative symptomatic urine frequently reveals species with uropathogenic potential, indicating that a negative SUC cannot reliably exclude infection (Heytens et al. 2017; Sathiananthamoorthy et al. 2019). Low-level bacteriuria, fastidious and anaerobic microbes, and polymicrobial infections are also routinely missed, yet the use of SUC for diagnosis drives empirical antibiotic prescribing (Bermudez et al. 2025).

It is now widely recognized that urine from an asymptomatic, presumably healthy, bladder is not sterile. Enhanced culture and DNA sequencing have shown the adult female urinary tract harbors a resident microbial community (called the urobiome) that SUC had failed to capture (Wolfe et al. 2012; Fouts et al. 2012; Khasriya et al. 2013; Hilt et al. 2014). Expanded quantitative urine culture (EQUC), which employs larger sample volumes, multiple media, and a wide range of atmospheric and incubation conditions, detects live microbes in the majority of specimens reported as “no growth” by SUC, and recovers a considerably broader range of species, including those considered to have uropathogenic and uroprotective potential (Hilt et al. 2014; Coorevits et al. 2016; Price et al. 2016; Deen et al. 2023). EQUC provides a richer, far more complete picture of the urobiome than SUC (Pearce et al. 2014; Thomas-White et al. 2018). Nonetheless, because it remains culture-based, EQUC retains inherent selection biases, cannot readily distinguish closely related species or strains, and is more labor- and time-intensive than SUC, limiting uptake as a routine clinical tool. As such, it has been recommended for use when SUC consistently returns a negative result (Price et al. 2016).

Culture-independent molecular methods, such as PCR-based assays, 16S rRNA gene profiling, and shotgun metagenomics, offer an increasingly attractive alternative. However, they all depend greatly on the efficient, unbiased extraction of microbial DNA.Yet extraction is not a neutral process, and incomplete or uneven cell lysis, particularly of hard-to-lyse Gram-positive organisms, systematically distorts apparent sample composition. Extraction methodology has repeatedly been found to be among the single largest sources of variation in microbiome analysis, frequently exceeding biases due to sequencing or bioinformatic workflows (Brooks et al. 2015; Costea et al. 2017; Kool et al. 2023).

Among sequencing approaches, short-read amplicon sequencing of the 16S rRNA gene has been the workhorse of microbiome research, allowing simultaneous, culture-free detection of many bacterial taxa from a single sample. However, it also suffers from limitations; confident taxonomic resolution is frequently restricted to the genus or even family level, results are relative rather than absolute, and reliance on PCR introduces amplification and primer biases that can distort community structure and generate false positives, particularly in low-biomass samples such as urine (Fouts et al. 2012; Greninger 2018; Kennedy et al. 2023). Because it analyzes only a short, single PCR amplicon, short-read 16S rRNA gene sequencing is limited to a small fraction of the available genomic information. In contrast, the more recently established long-read 16S rRNA gene sequencing improves taxonomic resolution by interrogating variable genomic segments in a longer amplicon but retains the amplification bias and relative quantitation inherent to any PCR-based approach (Jin, Liu, and Shiroguchi 2024; Johnson et al. 2019).

Shotgun metagenomic sequencing, which surveys total sample microbial DNA, offers greater taxonomic breadth and access to strain-level and functional information, but conventional short-read shotgun workflows are costly, computationally demanding, and constrained in urine by low microbial biomass and high host-DNA content (Wensel et al. 2022).

Applied clinically, metagenomic next-generation sequencing nonetheless can detect all DNA-based microbes (*i.e.*, bacteria, fungi and other eukaryotic microbes, and DNA viruses) in culture-negative and polymicrobial specimens and can shorten time to result, supporting its diagnostic value in UTI. It also can detect resistance determinants (Bermudez et al. 2025; Jeff et al. 2025). Long-read shotgun sequencing extends these gains, enabling reconstruction of high-quality metagenome-assembled genomes, including distinct *E. coli* subpopulations, directly from urine (Liu et al. 2026), but, like short-read versions, workflows can be both costly and resource-hungry.

A Nanopore-based long-read metagenomic sequencing workflow derivative, q3MetaG, was reported to mitigate many of these issues (Ferneyhough et al., 2025). It pairs robust cell-lysis and PCR-free library preparation with an internal calibration standard allowing direct reporting of absolute microbial load as cells per milliliter for each species identified with minimal extraction and processing bias. Applied as proof of principle to asymptomatic urogenital samples, the method reproduced established features of the healthy urinary and vaginal microbiomes while additionally resolving individual strains and quantifying absolute biomass across bacteria, fungi, viruses, plasmids and phages (Ferneyhough et al. 2025).

Critically, these technologies have rarely been evaluated head-to-head on the same specimens, leaving clinicians and researchers without a direct evidence base on which to judge their relative performance. To address this gap, we applied all five workflows: SUC, EQUC, short-read 16S rRNA gene sequencing, long-read 16S rRNA gene sequencing, and long-read shotgun (q3MetaG) metagenomic profiling to parallelized aliquots of a single cohort of urine samples. By comparing the microbes detected, the quantitative and taxonomic resolution achieved, and the clinical interpretation each method supports, we sought to define the relative strengths, limitations and clinical applicability of these approaches.

## METHODS

### Study design

This study is an exploratory analysis of the Peezy vs Clean Catch for Detecting Asymptomatic Bacteriuria (PDAB) randomized controlled trial; primary results were reported separately (Dhillon et al. 2026). Briefly, the PDAB study sought to determine if the Peezy® Midstream Urine Specimen Collection device could reduce the rate of urine culture contamination compared to standard midstream voided urine specimens in a population of pregnant patients being screened for asymptomatic bacteriuria. The Peezy® is a funnel-shaped device designed to discard approximately 10 mL of the initial voided urine before the remainder enters the sterile collection chamber (Southworth et al. 2019). This study was carried out at a single academic medical center, approved by the institutional review board (LU 216104) and registered on ClinicalTrials.gov (NCT 05554081). Patients were eligible at their initial prenatal visit if they were age >18 years, gestational age <20 weeks, and preferred English for medical communication. Patients with a history of recurrent UTI or ongoing antibiotic use were excluded. Participants’ data were collected from the medical chart. Their demographics have been published (Dhillon et al. 2026). Because the primary results indicated no significant difference in urobiome composition regardless of urine collection method (Dhillon et al. 2026), and because each participant functioned as their own control, urine samples from both groups of participants were combined into a single group for this exploratory study.

After collection, the urine specimens from each participant were divided into 3 samples: one for analysis in the hospital’s clinical laboratory, one for culture-based analysis in the translational researcher’s laboratory, and one for DNA-based analysis in the translational researcher’s laboratory. Samples were transported to the clinical laboratory according to standard medical practice. Some samples were maintained in a 4°C refrigerator for fewer than 4 hours before being transported to the translational researcher’s laboratory, where they were immediately subjected to culture-based detection methods or divided into two aliquots for subsequent sequencing. To each of these aliquots, we added a nucleic acid preservative (AssayAssure®, Sierra Molecular Corp, Princeton, NJ, LOT G3030C) in a 1:9 ratio, before freezing at -80°C. One aliquot was kept for 16S rRNA gene sequencing (short-read and long-read). The other aliquot was shipped on dry ice to the Systems Biology Laboratory (Abingdon, UK) for long-read shotgun metagenomic sequencing. Only samples transported to the translational researcher’s laboratory were considered in the analyses reported here.

The purpose of the current study was to identify the strengths and weaknesses of several different bacterial detection methods: two culture-based methods (SUC and EQUC) and three DNA-based methods (short-read 16S rRNA gene sequencing, long-read 16S rRNA gene sequencing, and shotgun metagenomic sequencing). For an overview of the study design, see **Figure 1A**.

**Figure 1.**
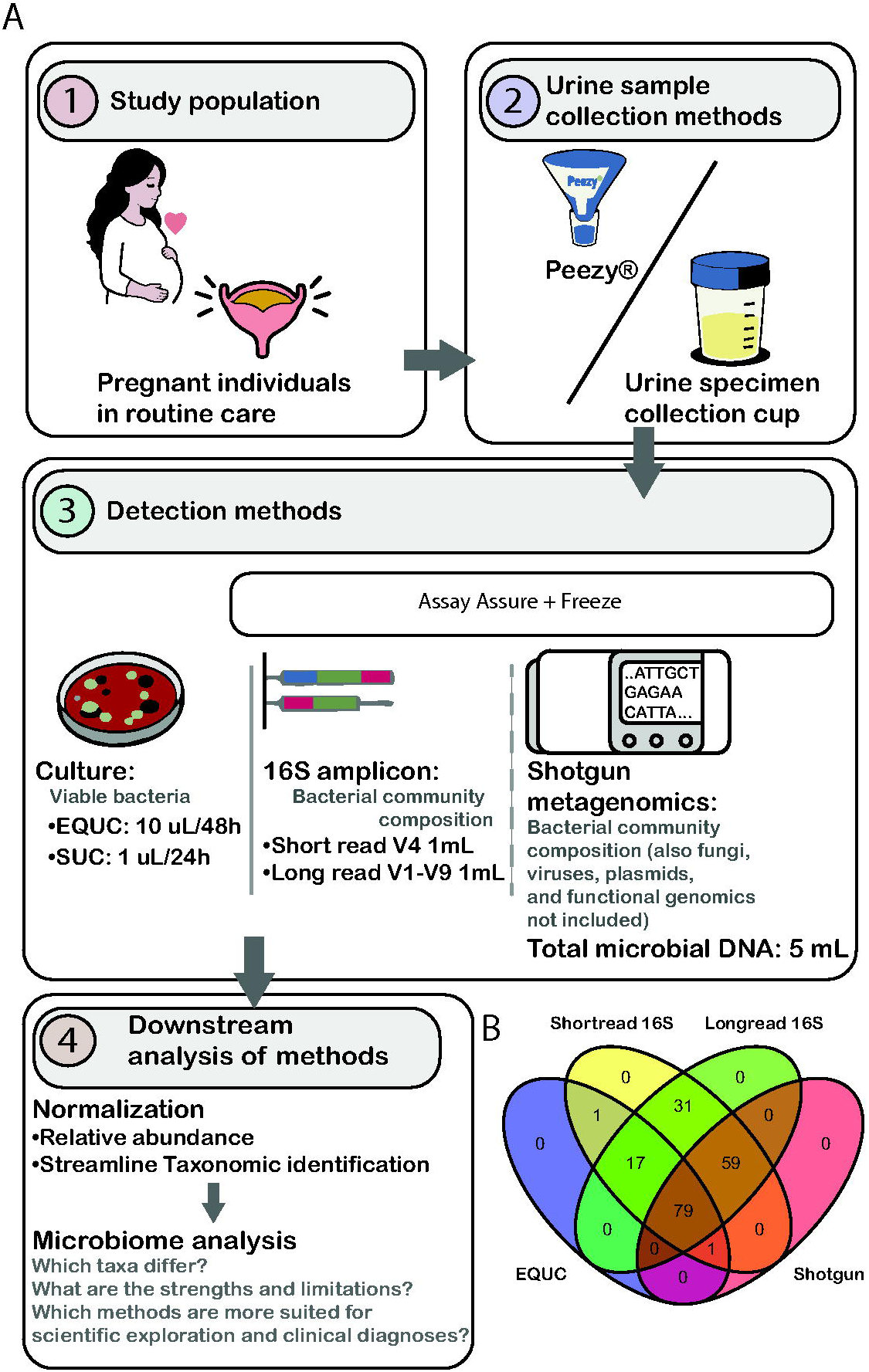
**Panel A.** Experimental Design. **Panel B.** Venn diagram portraying the intersection of positive results for 4 detection methods: EQUC, short-read and long-read 16S rRNA gene sequencing, and long-read shotgun metagenomic sequencing.

### Culture-based detection methods

Techniques for SUC and EQUC have been reported previously (Hilt et al. 2014; Price et al. 2016). Briefly, SUC involves inoculation of 1 μL of urine onto one MacConkey agar plate and onto one Blood Agar Plate (BAP), with incubation in ambient atmosphere at 35°C for 24 hours. The detection level was 1,000 colony-forming units (CFU)/mL, represented by 1 colony of growth on either plate. In contrast, EQUC involves inoculation of 10 μL of urine onto multiple agar plates (BAP and Colistin Nalidixic Acid [CNA, BD BBL™ Prepared Plated Media]) with incubation in 5% CO_2_ at 35°C for 48 hr, and onto CDC Anaerobe 5% sheep blood agar plates (BD BBL™ Prepared Plated Media), and incubation under anaerobic conditions at 35°C for 48 hr. The detection level was 100 CFU/mL, represented by 1 colony of growth on any of the plates. Colonies were identified by Matrix-Assisted Laser Desorption/Ionization Time-of-Flight (MALDI-ToF). Contrary to typical clinical microbiology laboratory reporting standards, all detected taxa were reported.

### For 16S rRNA gene sequencing: genome extraction and library preparation

For both short-read and long-read 16S rRNA gene sequencing, genomic DNA was extracted from 1 mL of urine using the following previously validated protocols. In short, enzymatic treatment, consisting of 4 mg Lysozyme, 3 kU Mutanolysin, and 4 μg Lysostaphin in buffer (20 mM Tris, 2 mM EDTA, 1.2% Triton, pH 8.0), was added to each sample and incubated at 37°C for 5 hours, shaking intermittently at 1000 rpm (Eppendorf Thermomixer with SmartBlock DWP1000). Samples were then processed using the DNeasy 96 Blood & Tissue kit (Qiagen, Germantown, MD) according to the manufacturer’s instructions. To assess potential DNA contamination, extraction negative controls (no urine) were processed with the samples and sequenced in every run.

The urine samples were sequenced by SeqCenter (Pittsburgh, PA). For short-read 16S rRNA gene sequencing, they were amplified using Zymo Research’s Quick-16S™ kit with phased primers targeting the V4 region of the 16S rRNA gene (5’-GTGYCAGCMGCCGCGGTAA-3’, 5’-GGACTACNVGGGTWTCTAAT-3’). Following clean-up and normalization, samples were sequenced on a P2-300cyc NextSeq2000 Flowcell to generate 2x151bp paired end (PE) reads. For long-read 16S rRNA gene sequencing, they were amplified via 1-step PCR using Zymo Research’s Quick-16S™ Full-length Library Kit with primers targeting V1-V9 regions of the 16S rRNA gene (27f; 5’-AGRGTTYGATYMTGGCTCAG-3’; 1492r; 5’-RGYTACCTTGTTACGACTT-3’). A MagBead cleanup was performed on the library pooled by volume and then prepared for sequencing using the PacBio HiFi prep kit 96 with the SMRTbell® barcoded adapter plate 3.01. Final libraries underwent the PacBio binding and cleanup steps using the Revio SPRQ Polymerase kit. Prepared libraries were sequenced on the PacBio Revio with a 12hr run time.

### For 16S rRNA gene sequencing: data processing, decontamination, and taxonomic identification

For short-read and long-read 16S rRNA gene sequencing, adaptors were removed, the raw reads assessed for quality, and filtered using FastQC (v. 0.11.9) and Cutadapt (v. 3.7) (Martin 2011). The DADA2 pipeline was used for read merging, chimera checking, and ASV identification (Callahan et al. 2016). To achieve taxon identification, BLCA (Gao et al. 2017) was used to query the NCBI and Greengenes2 databases (McDonald et al. 2024). For each ASV, identification with confidence at the lowest taxonomic level was used for downstream analyses. ASVs with taxonomic confidence scores below 69.5% were classified as "unclassified/unknown." ASVs assigned to the genus *Lysobacter* were removed, as this genus is a known contaminant associated with the enzyme achromopeptidase used in the DNA extraction procedure. Low-abundance ASVs were removed. Then, contaminant ASVs were determined via a scoring method that encompasses results from Decontam (Davis et al. 2018), nonparametric statistical testing (Kruskal-Wallis p≥0.5), mean comparison (*i.e.*, mean counts of samples were less than mean counts of extraction controls), and whether sample read counts were less than 5x greater than extraction controls, as described previously (Gevelinger et al. 2024).

### For long-read shotgun metagenomic sequencing: genome extraction and library preparation

For each sample, 2 mL of frozen urine was used as the input. Samples were thawed and warmed (37 °C, 30 min) to prevent co-precipitation of lipids and proteins, and cells were recovered by centrifugation (14,000 × g, 5 min). Host cells were selectively lysed and free host DNA removed by enzymatic host depletion (ZymoBIOMICS HostZero chemistry: host-depletion solution, followed by microbial selection buffer with nuclease digestion at 37 °C and subsequent proteinase K inactivation). The retained microbial fraction was spiked with a fixed titer of an internal calibration standard (ZymoBIOMICS Spike-in Control) to enable absolute quantification, then lysed by heating (95 °C, 5 min) and mechanical bead-beating (FastPrep, MP Biomedicals), and sheared by sonication (Bioruptor, Diagenode). Microbial DNA was captured on a silica spin column, washed, eluted in nuclease-free water, and quantified by fluorometry (Quant-iT PicoGreen/AccuBlue, read on a BMG PHERAStar against a DNA standard curve). Sample tracking, reagent lot control, and yield calculation were managed within the laboratory information management system (LIMS).

Sequencing libraries were prepared from extracted DNA (target ∼100 ng; maximum 400 ng input) using the Oxford Nanopore ligation-based native barcoding kit (SQK-NBD114.24). DNA was end-repaired and A-tailed (NEBNext FFPE DNA Repair and NEBNext Ultra II End Prep), purified with AMPure XP beads (Beckman Coulter), ligated to native barcodes and then to sequencing adapters, with bead-based clean-up and size selection between steps. Barcoded libraries were quantified, pooled, and eluted ready for loading. Library preparation was performed manually or by an equivalent semi-automated protocol on a Tecan Fluent liquid-handling workstation.

Pooled libraries were loaded onto R10.4.1 flow cells and sequenced on a GridION device (Oxford Nanopore Technologies) under MinKNOW control, typically as overnight runs (∼14 h). Basecalling was performed with the current Oxford Nanopore basecaller (Dorado).

#### For long-read shotgun metagenomic sequencing: data processing, decontamination, and taxonomic identification

Raw Oxford Nanopore sequencing reads were initially assessed using NanoStat v1.6.0 (De Coster et al. 2018) to summarize read length and quality distributions. Reads were subsequently filtered with NanoFilt v2.8.0 (De Coster et al. 2018), retaining sequences with a minimum Phred quality score of 12 (Q ≥ 12). No length-based filtering was applied.

To remove host and non-target sequences, quality-filtered reads were sequentially aligned against the reference genomes of the sequencing kit-associated bacterial contaminants *Allobacillus halotolerans* strain BCRC 17939 (RefSeq accession GCF_028743495.1) and *Imtechella halotolerans* strain JCM 17677 (RefSeq accession GCF_028743515.2), followed by alignment against the human reference genome GRCh38 (RefSeq accession GCF_000001405.39) to remove host-derived reads, as dictated by our approved IRB protocol. Alignments were performed using minimap2 v2.24 (r1122) (Li 2018) with the Oxford Nanopore preset (-x map-ont) and secondary alignments disabled (--secondary=no). Alignment files were sorted using SAMtools (Danecek et al. 2021), and only reads that did not map to any reference sequence were retained for downstream metagenomic analyses.

Taxonomic profiling was performed using Kraken2 v2.1.3 (Wood, Lu, and Langmead 2019) with a custom reference database. Species-level abundance estimates were refined using Bracken v3.0.1(Lu et al. 2017), assuming a read length of 1,000 bp. Taxa with fewer than 10 estimated reads were excluded, and only taxa meeting this threshold were retained for downstream analyses.

#### Bioinformatic and biostatistical analysis

To fairly describe the bacterial distribution among the five different methods, steps were taking to normalize their datasets to one another.

First, culture data (SUC and EQUC) contained CFU/ml of species identified by MALDI-ToF. Species information was used to determine taxonomic lineage using the taxize R package (v.0.10.1) (Chamberlain and Szocs 2013), which sourced taxonomy information from the NCBI database. For 16S rRNA gene profiling (short-read 16S V4 region and long-read 16S rRNA V1-V9 region) data contained read count information along with taxonomy. Taxonomy lineage at the genus or family level were assessed to maintain consistency, using the taxize R package (Chamberlain and Szocs 2013).

Next, we determined which of the 188 participants had their urine sample assessed by all methods. Of 188 samples, 101 samples were cultured, with 98 EQUC-positive, and 85 SUC-positive. Short-read 16S rRNA sequencing resulted in 188 positive samples, while long-read 16S rRNA sequencing resulted in 186 positive samples. All 139 samples that were subjected to long-read shotgun metagenomic profiling was positive. Seventy-nine participants were positive by EQUC, short-read 16S rRNA gene sequencing, long-read 16S rRNA gene sequencing, and long-read shotgun sequencing. This intersection of method-positive participants was compared as follows.

Alpha (within sample) diversity and beta (between sample) diversity were computed using statistical packages phyloseq (McMurdie and Holmes 2012) and vegan (Jari Oksanen and B. O’Hara 2015) in R (R Foundation for Statistical Computing, Vienna, Austria). Alpha diversity was used to assess bacterial populations within each participant sample, using the following indices: Observed, Chao1, and Ace as measures of richness; Pielou as a measure of evenness; Shannon and Simpson as measures of overall diversity. Each index was calculated at the family, genus and species levels. Statistical significance was calculated with pairwise tests with p-values ≤0.05 considered statistically significant.

Because culture and sequencing methods output counts (*i.e.*, CFU/mL or read counts) are unique to their respective methodology, raw counts were converted to relative abundances and prevalence, and then aggregated at the genus and species levels, in order to be presented in an unbiased manner. We then compared the 5 methodologies calculating beta diversity using both Bray-Curtis Dissimilarity and Canberra Distance indices ordinated by Principal Coordinate Analysis (PCoA). Statistical significance was determined by Permutational Multivariate Analysis of Variance (PERMANOVA) with p-values ≤0.05 considered statistically significant.

Cluster analysis of the 5 methodologies were performed such that the relative abundances were converted to prevalence (with no threshold restrictions, relative abundance >0.0 were permitted) at the genus and species levels, Bray-Curtis dissimilarity indices were determined and were then clustered by way of UPGMA method (via RStats).

Differential abundance analysis (DAA) using ANCOMBC function at species level. Differential abundance analysis (DAA) was performed by analysis of compositions of microbiomes with bias correction (ANCOMBC) (Lin and Peddada 2020). The statistical approach includes normalization of read counts using a process similar to log-ratio transformations along with controlling false discovery rates. Raw genera counts were used as input for the “ANCOMBC” function, and adjusted p values (<0.05) were set to Benjamini-Hochberg.

### Data accessibility

The raw sequencing data can be accessed under BioProject ID: PRJNA1465356, entitled “Detection methods of asymptomatic bacteriuria in an obstetric population.”

## RESULTS

### Study design

We previously published the results of a randomized controlled trial of the Peezy® Midstream Urine Specimen Collection Device compared to the conventional midstream voided “clean catch” method in an obstetrics population (Dhillon et al. 2026). Because we observed no significant difference between the two collection methods, all participant samples were considered in this exploratory analysis, which compared multiple methods of bacterial detection, including the SUC method typically used by clinical microbiology laboratories, an enhanced culture method (EQUC), short-read 16S rRNA gene sequencing targeting the V4 region, long-read near full-length (V1-V9) 16S rRNA gene sequencing, and long-read shotgun metagenomic sequencing (**Figure 1A**).

### SUC versus EQUC

Multiple reports have provided evidence that enhanced methods (*e.g.,* EQUC) outperform SUC (Price et al. 2016; Hochstedler et al. 2022; Deen et al. 2023; Khasriya et al. 2013; Coorevits et al. 2016). The results of our current analysis support this conclusion. Of the 188 participants who provided a urine sample, 101 were cultured. Of these 98, 98 (100.0%) were EQUC-positive, whereas only 85 (86.7%) were SUC-positive (**Figure 2**). More importantly, EQUC detected considerably more species (n=102) than did SUC (n=41) (**Table 1**).

**Figure 2.**
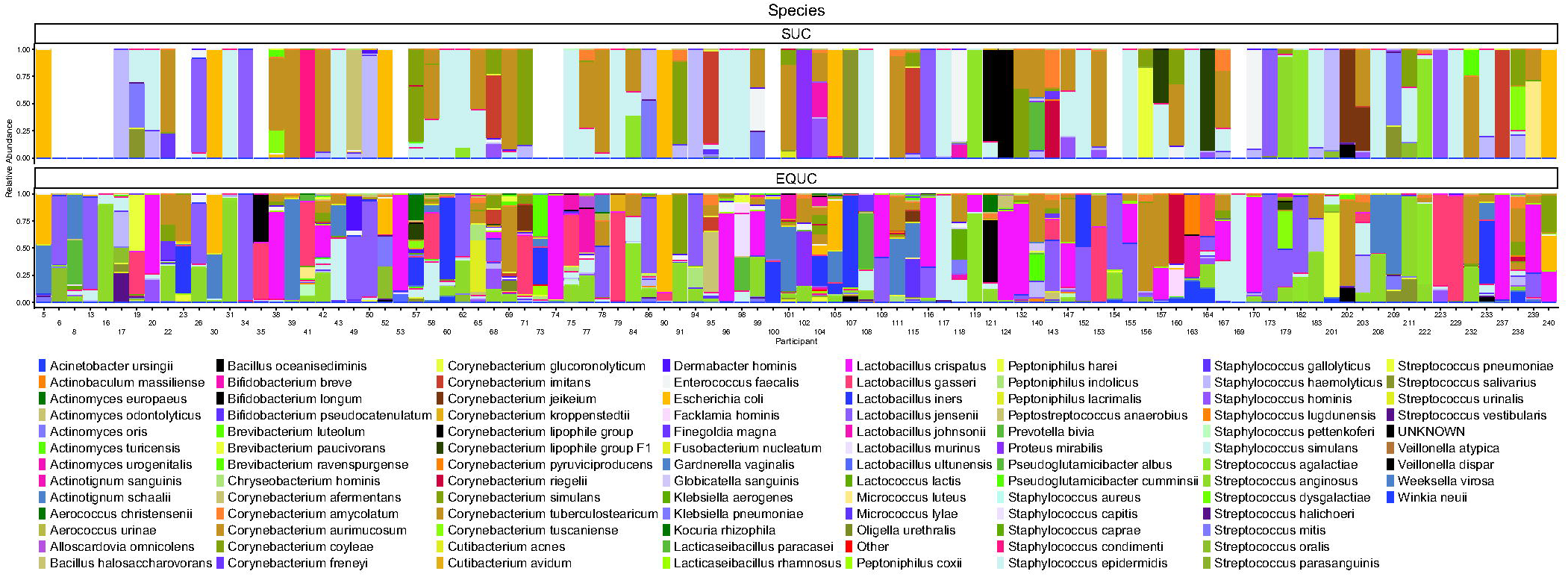
Comparison of SUC and EQUC results for all 101 cultured samples.

**Table 1.**
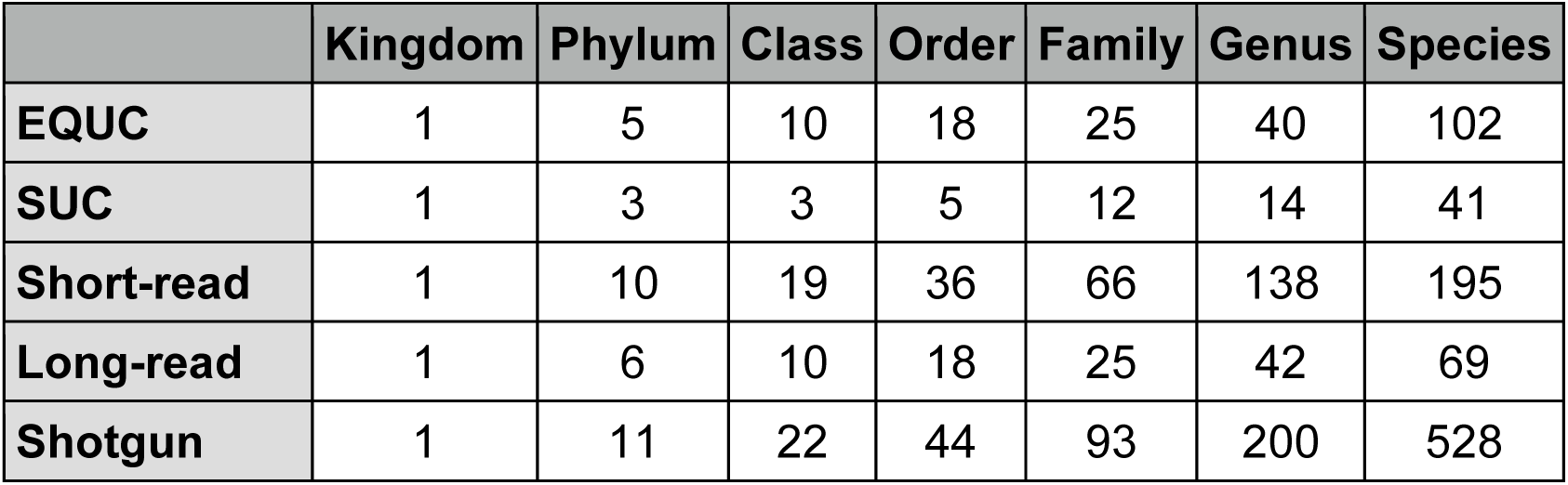
Number of unique taxa identified by each detection method.

### Sequencing versus culture methods

We next compared the number of samples that were positive by the 3 different DNA sequencing methods to those that were positive by EQUC. Whereas 98 participants were EQUC-positive (**Figure 1B**) with 102 species detected (**Table 1)**, 188 were positive by short-read 16S rRNA gene sequencing (195 species detected), 186 were positive by long-read near full-length 16S rRNA gene sequencing (69 species detected), and all 139 assessed by long-read shotgun sequencing were positive (528 species detected). To compare taxonomic identification across all detection methods, all subsequent analyses were performed on only the 79 samples that were positive by all 4 methods. For these 79 samples, we also compared the SUC results, whether they were positive or negative.

First, we compared taxonomic identification of ASVs obtained by either short-read or long-read 16S rRNA gene sequencing by querying both the NCBI and the Greengenes2 databases or querying either database alone (**Table 2**). When querying both databases, the number of reads with a confident identification increased at both the genus and species levels for both short- and long-read 16S rRNA gene sequencing methods. Long-read 16S rRNA gene sequencing, however, yielded a higher percentage of confident identifications (98% at the genus level, 88% at the species relative to 90% and 68%), especially at the species level.

**Table 2.** Taxonomic identification at the genus and species level querying either the Greengenes 2 (GG2) or the NCBI databases or both (MERGED) for both short-read (top) and near full-length long-read (bottom) 16S rRNA gene sequencing.

| **Short-read** <sup>a</sup> |  |  |  |  |
|  | **Genus** |  | **Species** |  |
|---|---|---|---|---|
| **GG2** | 1553 | 76% | 1098 | 53% |
| **NCBI** | 1754 | 85% | 1099 | 53% |
| **MERGED** | 1854 | 90% | 1398 | 68% |
| **Long-read** <sup>b</sup> |  |  |  |  |
|  | **Genus** |  | **Species** |  |
| **GG2** | 5085 | 82% | 4051 | 65% |
| **NCBI** | 5699 | 92% | 5045 | 81% |
| **MERGED** | 6086 | 98% | 5488 | 88% |

<sup>a</sup> Out of a total of 2055 ASVs.
<sup>b</sup> Out of a total of 6225 ASVs.

Next, we performed alpha (within sample) diversity analyses. At both the genus (**Figure 3A**) and species (**Figure 3B**) levels, SUC was the least rich (Observed, Chao1), the most even (Pielou), and the least diverse overall (Shannon, Simpson). In contrast, short-read 16S rRNA gene sequencing and shotgun metagenomic sequencing were the richest, but, along with long-read near full-length 16S rRNA gene sequencing, were also less even, especially short-read 16S rRNA gene sequencing.

**Figure 3.**
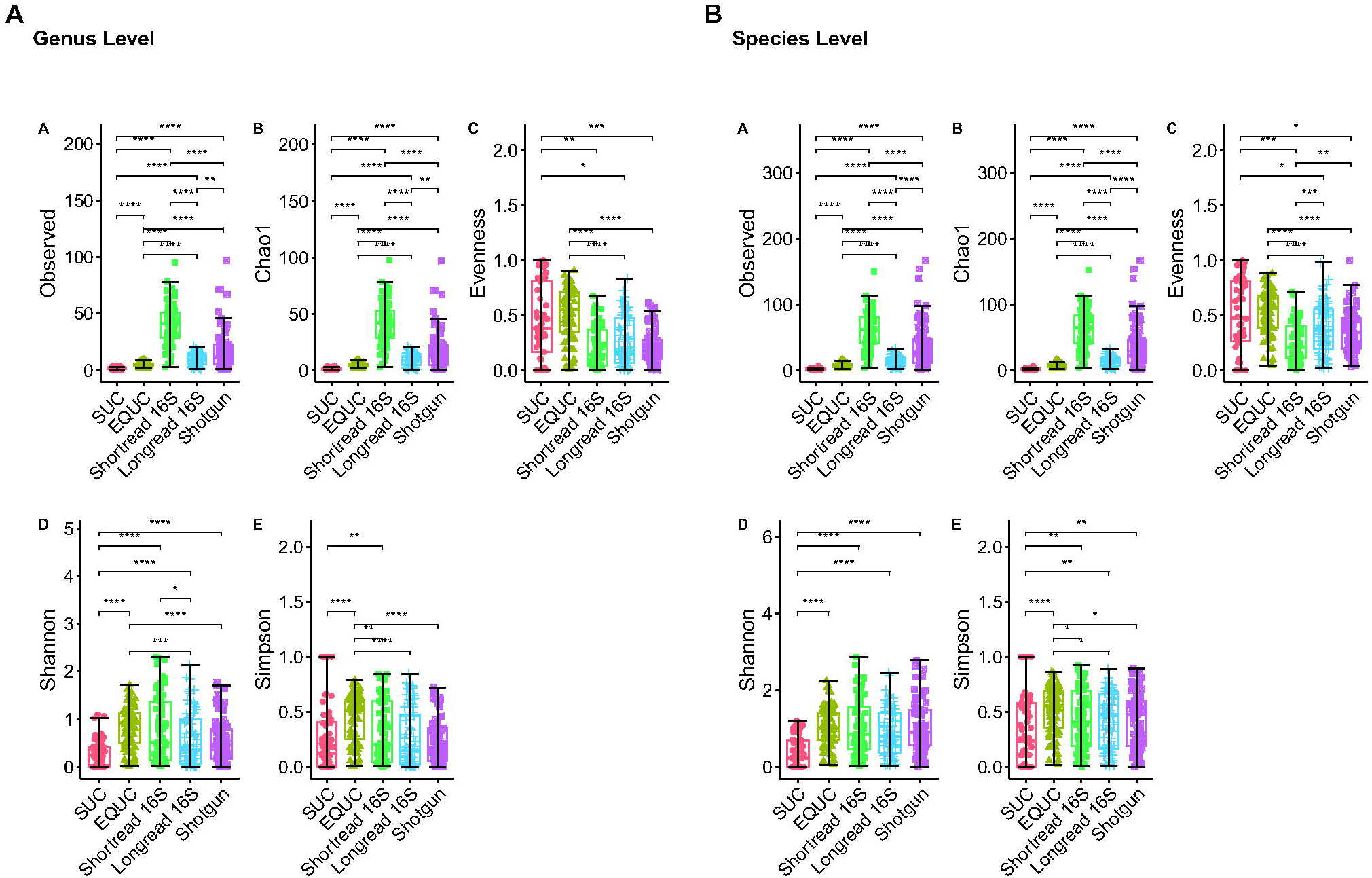
Alpha diversity analysis comparing all 5 detection methods. p = *<0.05, **<0.01, ***<0.001). **Panels A-E:** Genus level, **Panels F-J**: Species level.

Hierarchical cluster analysis showed, at both the genus and species levels, that the results of the three sequencing-based methods resemble each other more than they do the results of the two culture-based methods (**Figure 4A & B**). At the genus level, the results of the long-read methods were more similar to each other than those of the short-read method. At the species level, however, the results of the two 16S rRNA gene sequencing methods resembled each other more than they did the results of shotgun metagenomic sequencing.

**Figure 4.**
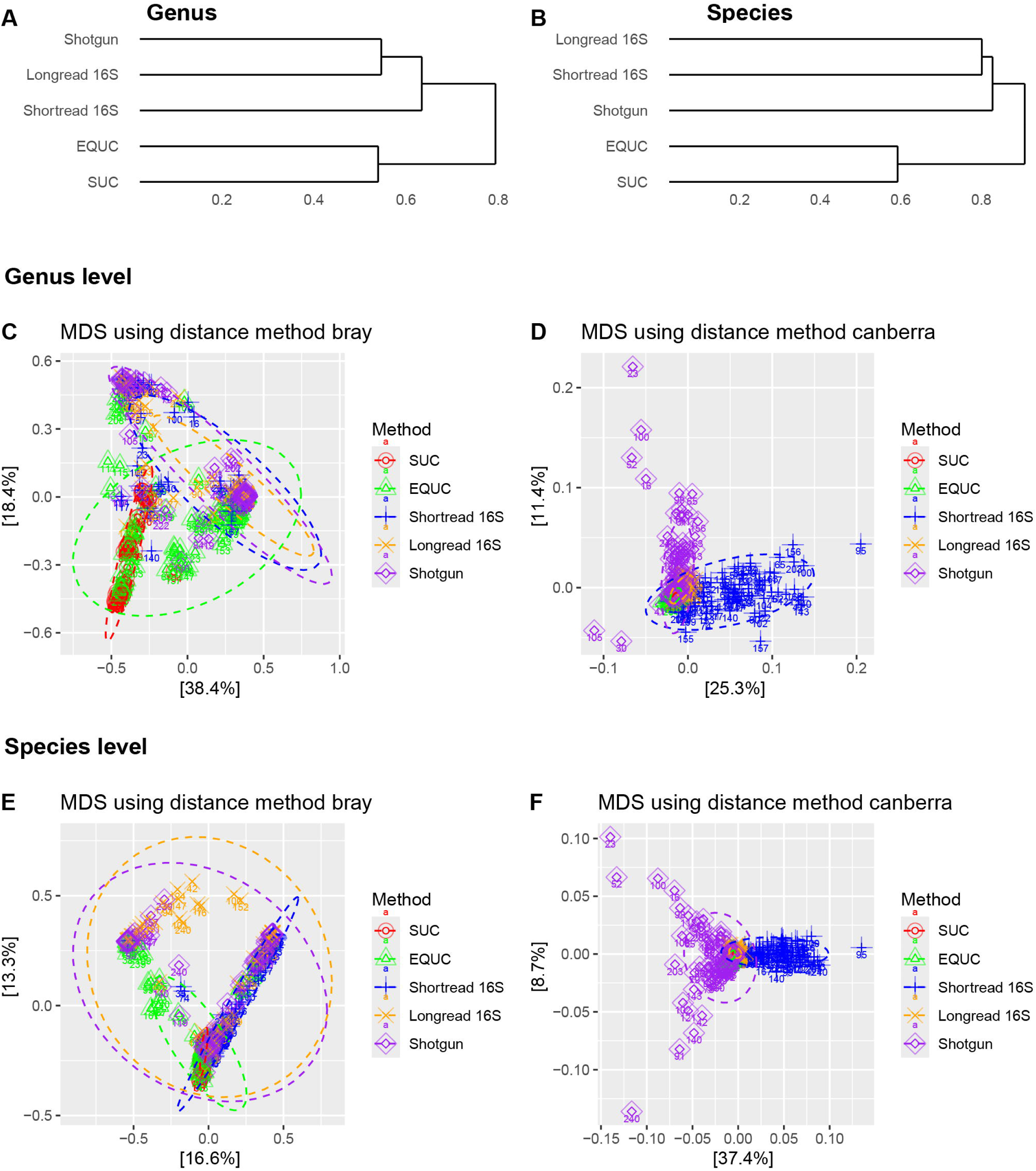
Hierarchical cluster analysis and beta diversity analyses comparing the 5 detection methods. **Panels A and B:** hierarchical clustering at the genus and species levels, respectively. **Panels C and D:** for genus level identification, beta diversity analysis using Bray-Curtis Dissimilarity and Canberra Distance indices, respectively, ordinated by Principal Coordinate Analysis (PCoA), a multidimensional scaling (MDS) method. **Panels E and F:** for species level identification, beta diversity analysis using Bray-Curtis Dissimilarity and Canberra Distance indices, respectively, ordinated by PCoA.

These results were supported by beta (between sample) diversity analysis calculated with both the Bray-Curtis Dissimilarity Index and the Canberra Distance Metric and ordinated by Principal Coordinate Analysis (PCoA) (**Figure 4C-F**). Bray-Curtis emphasizes predominant taxa, whilst Canberra is sensitive to rare taxa. At the genus level, PCoA ordination of Bray-Curtis distances showed the results of (1) the three sequencing approaches cluster together, indicating that they similarly identify the most predominant genera, (2) EQUC forms a unique cluster that partially overlaps with the sequencing methods, indicating that it misses some of the most predominant genera, and (3) SUC forms a subset of EQUC, indicating that this method fails to identify many of the genera identified by EQUC (**Figure 4C**). In contrast, PCoA ordination of Canberra distances shows that the short-read 16S rRNA gene and long-read shotgun metagenomic sequencing methods form distinct clusters from the other methods and from each other, indicating that these two methods uniquely identify more rare genera (**Figure 4D**). At the species level, Bray-Curtis distances revealed (1) greater similarity between the two long-read sequencing methods, (2) that EQUC and short-read 16S rRNA gene sequencing detect a subset of species identified by the two long-read sequencing methods, and (3) that SUC detects only a subset of species identified by the other four detection methods (**Figure 4E**). Once again, Canberra distances showed that long-read shotgun metagenomic and short-read 16S rRNA gene sequencing methods form distinct clusters from the other methods and from each other (**Figure 4F**).

Taken together, these results reveal that the long-read sequencing methods identify the most predominant species, the short-read and shotgun methods identify more rare species, and SUC identifies a small subset of species identified by the other methods, including EQUC.

To show the differences between the different detection methods, we constructed histograms showing the top 92 genera and 98 species ordered in parallel by participant number (**Figure 5A & B**). At the genus level, it appears that all three sequencing methods generally yielded similar results; for example, when *Escherichia* was identified by one method, the other two also identified *Escherichia* (gold). This also was generally true for *Lactobacillus* (blue) and *Gardnerella* (maroon). However, at the species level, EQUC often outperformed SUC, identifying species considered to have pathogenic potential, including *E. coli*, *E. faecalis, Gardnerella vaginalis, Klebsiella pneumoniae, Staphylococcus aureus, S. agalactiae,* and *S. anginosus* when SUC did not (**Table 3**). EQUC also did a better job of identifying species generally considered to be protective, including *L. crispatus, gasseri, iners, jensenii, mulieris,* and *paragasseri*. In some cases, both long-read methods (16S rRNA gene sequencing and shotgun sequencing) outperformed short-read 16S rRNA gene sequencing. Whereas the long-read methods identified *E. faecalis* at the species level, short-read 16S rRNA gene sequencing could only identify it at the genus level. With the exception of *L. iners,* this was also generally true of the *Lactobacillus* species. In contrast, for *S. agalactiae* and *S. anginosus*, the short-read method did about as well as the long-read methods, and both EQUC and all three sequencing methods outperformed SUC. The long-read shotgun metagenomic sequence method did well to identify most *Lactobacillus* species, but it was particularly adept at distinguishing the various species of *Gardnerella*. For the complete list of taxa detected by any of the detection methods, see **Supplemental Tables 1 and 2**.

**Figure 5.**
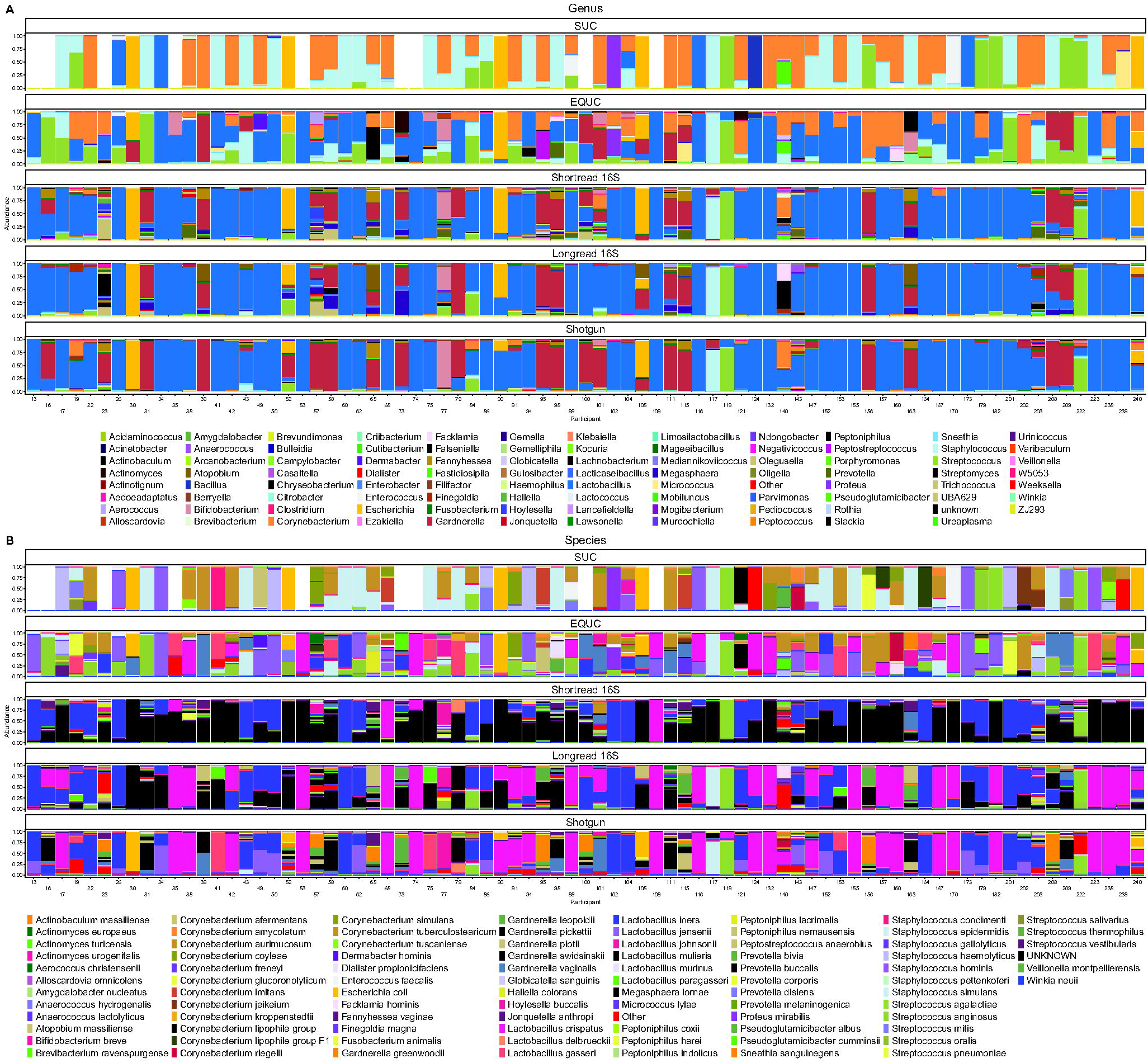
Histograms comparing all 5 detection methods. **Panel A:** genus level; **Panel B:** species level.

**Table 3.** The ability of the 5 detection methods to identify select taxa^a^.

| Taxon | SUC | EQUC | Short-read | Long-read | Shotgun |
|---|---|---|---|---|---|
| *Aerococcus* species | 0 | 0 | 24 | 0 | 0 |
| *A. christensenii* | 0 | 2 | 37 | 17 | 45 |
| *A. urinae* | 0 | 5 | 26 | 0 | 6 |
| *Escherichia* species | 0 | 0 | 35 | 10 | 0 |
| *E. coli* | 8 | 13 | 0 | 0 | 15 |
| *Enterococcus* species | 0 | 0 | 29 | 0 | 0 |
| *E. faecalis* | 6 | 20 | 0 | 16 | 13 |
| *Klebsiella pneumoniae* | 1 | 3 | 0 | 0 | 1 |
| *Proteus mirabilis* | 1 | 1 | 1 | 0 | 2 |
| *Serratia marcescens* | 0 | 0 | 0 | 0 | 2 |
| *Staphylococcus aureus* | 1 | 7 | 0 | 0 | 1 |
| *Staphylococcus saprophyticus* | 0 | 1 | 0 | 0 | 1 |
| *Streptococcus agalactiae* (GBS) | 6 | 10 | 15 | 12 | 24 |
| *Streptococcus anginosus* | 1 | 37 | 58 | 47 | 27 |
| *Lactobacillus* species | 0 | 0 | 65 | 21 | 0 |
| *L. crispatus* | 0 | 19 | 7 | 64 | 38 |
| *L. gasseri* | 0 | 12 | 0 | 1 | 35 |
| *L. iners* | 4 | 21 | 0 | 0 | 31 |
| *L. jensenii* | 0 | 8 | 73 | 79 | 58 |
| *L. mulieris* | 0 | 0 | 0 | 31 | 0 |
| *L. paragasseri* | 0 | 0 | 0 | 18 | 0 |
| *Gardnerella* species | 0 | 0 | 45 | 27 | 0 |
| *G. greenwoodii* | 0 | 0 | 0 | 0 | 25 |
| *G. leopoldii* | 0 | 0 | 23 | 23 | 33 |
| *G. pickettii* | 0 | 0 | 0 | 0 | 29 |
| *G. piotii* | 0 | 0 | 0 | 0 | 33 |
| *G. swidsinskii* | 0 | 0 | 0 | 25 | 38 |
| *G. vaginalis* | 0 | 13 | 43 | 24 | 39 |

<sup>a</sup> In only the 79 participants who were positive for EQUC and the three sequencing methods.

The previous analysis was based upon absolute enumeration (*i.e.,* CFU/mL or read counts). Therefore, we performed differential abundance analysis (DAA) by analyzing compositions of microbiomes with bias correction (ANCOMBC), which uses normalized data to identify taxa that differed significantly between detection methods (**Figure 6**). Relative to SUC, EQUC was more adept at detecting *E. faecalis, G. vaginalis, Lactobacillus* species (*crispatus, gasseri, iners, & jensenii*), and *S. anginosus.* Both SUC and EQUC were better able to detect *Corynebacterium* species (*amycolatum* & *auricumosum*), *E. faecalis*, and *Staphylococcus* species (*epidermidis, haemolyticus,* & *hominis*) than any of the sequencing methods. All three sequencing methods were more adept at detecting *Aerococcus christensenii*, *Finegoldia magna,* and *Fannyhessea vaginae/Atopobium massiliense.* Both16S rRNA gene sequencing methods were better at detecting *Escherichia* species. Both EQUC and long-read shotgun sequencing did better than both the 16S rRNA gene sequencing methods at identifying *G. vaginalis* and distinguishing the various *Lactobacillus* species. The long-read methods (16S rRNA gene and shotgun) were best at distinguishing *Gardnerella* species. Short-read 16S rRNA gene sequencing was most adept at identifying *Megasphaera lornae* and *Prevotella* species (*bivia* & *buccalis*).

**Figure 6.**
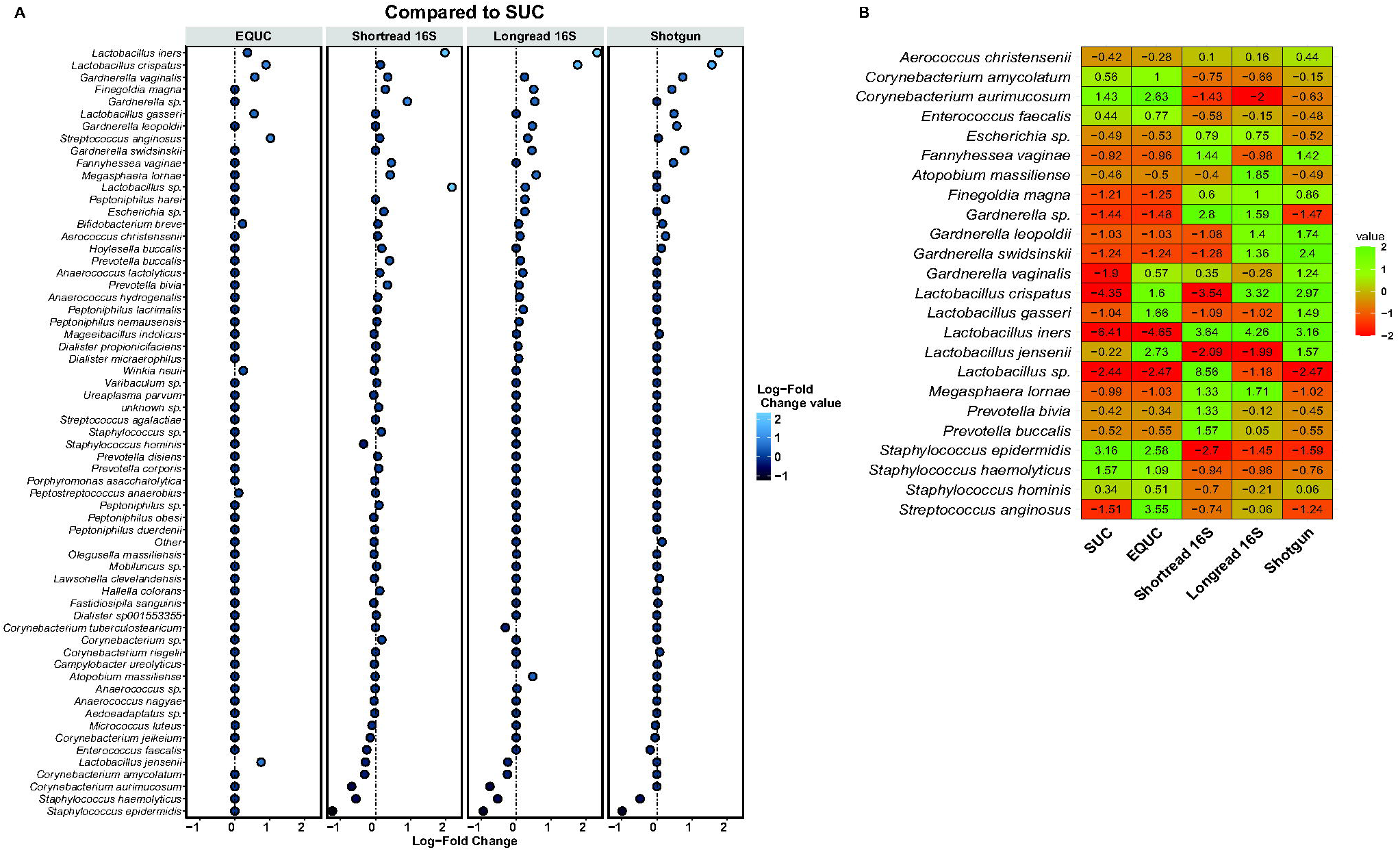
ANCOM-BC analysis. **Panel A.** Results comparing each method against the other 4 methods. **Panel B.** Summary of ANCOM-BC results.

## DISCUSSION

### Summary of the Results

This study demonstrates that the answer to a seemingly simple question — what is in this urine sample? — depends substantially on how the urine is analyzed. Using parallel aliquots from the same participant specimens, we compared two culture-based approaches, standard urine culture (SUC) and expanded quantitative urine culture (EQUC), with three sequencing-based approaches, short-read 16S rRNA gene sequencing, long-read 16S rRNA gene sequencing, and long-read shotgun metagenomic sequencing. Across these five methods, we observed marked differences in positivity, species recovery, richness, taxonomic resolution, and overall representation of the urinary microbial community. The central finding is not merely that some methods detect more than others, but that different methods generate meaningfully different biological pictures from the same specimen.

Among the culture-based methods, EQUC outperformed SUC in both positivity and species recovery. Of the 98 cultured samples, 98 were positive by EQUC and 85 were positive by SUC, and EQUC recovered a broader set of microbes, including taxa considered uropathogenic as well as microbes often associated with a *Lactobacillus*-predominant urogenital state. These findings are consistent with prior work showing that SUC either underestimates or does not detect bacterial taxa that can be recovered under enhanced culture conditions (Brubaker et al. 2023; Price et al. 2016; Moreland et al. 2024). Even within culture-based diagnostics, therefore, methodological choice shaped the resulting biological interpretation.

The sequencing-based methods broadened detection further, but in different ways. The short-read 16S method was positive for all 188 samples, the long-read method for 186 samples, and long-read shotgun metagenomics for all 139 samples assessed by that method. The short-read 16S and long-read shotgun metagenomic methods recovered high richness, whereas the long-read 16S method improved confident taxonomic assignments, particularly at the species level, relative to the short-read method. In addition, the sequencing-based methods more closely resembled one another than they resembled the culture-based methods, indicating that culture and sequencing generate overlapping but distinct views of urinary microbial composition.

Taxon-level findings reinforce this point. Relative to SUC, EQUC improved recovery of many species, including *E. coli*, *E. faecalis*, *A. christensenii*, *C. aurimucosum*, *F. magna*, and multiple *Lactobacillus* species. The long-read methods better distinguished certain taxa at the species level, including *Gardnerella* species, whereas the short-read 16S method appeared particularly effective for detection of some lower-abundance or fastidious taxa, including *M. lornae* and certain *Prevotella* species. Long-read shotgun metagenomics and EQUC also outperformed the 16S-based methods for some microbes, including *G. vaginalis*. Taken together, these findings show that the five workflows cannot be arranged along a single performance gradient. Rather, each has areas of relative strength, and each emphasizes different properties of the specimen.

Importantly, no single method was uniformly optimal across all analytical objectives. Instead, each approach contributed a distinct view of the urinary microbial community. This is the dominant governing insight from the study and should shape both future research design and any attempt to translate these methods into clinical use.

#### Research Implications

The research implications of these findings are immediate. For urobiome studies, method selection should be driven by the specific biological question being asked, because the five methods evaluated here do not provide interchangeable information. Different methods optimized different endpoints—viable microbe recovery, broad community profiling, species-level taxonomic resolution, rare-taxon detection, or potentially calibrated abundance—and those endpoints should not be collapsed into a single undifferentiated notion of performance.

If the research goal is to recover living microbes for downstream experimentation, EQUC is the stronger culture-based option. Because it identified more positive samples and recovered more species than SUC, EQUC provides a broader viable snapshot of the urine specimen while preserving the ability to culture isolates. This matters for work involving antimicrobial susceptibility testing, strain preservation, co-culture experiments, genome sequencing of isolates, and mechanistic studies of host–microbe interaction. By contrast, SUC appears insufficient as a stand-alone method for comprehensive microbiome-oriented studies, because it underestimates both positivity and taxonomic diversity. As with any culture-based workflow, microbe identification also may require downstream taxonomic identification methods such as MALDI-TOF, biochemical testing, or whole genome sequencing; this can be an advantage when viable isolates are needed, but it adds procedural dependence relative to direct sequencing-based readouts.

If the goal is broad community profiling across many samples, short-read 16S rRNA sequencing remains useful. In this study, it produced nearly universal positivity and high richness, making it practical for ecological analyses, cohort comparisons, and hypothesis-generating work. It is likely to remain attractive where funding is limited or the study is exploratory—for example, in pilot projects designed to investigate some unstudied or understudied problem. Its lower cost relative to more information-rich methods is a practical inference rather than a conclusion established by this dataset, but it is an important real-world consideration in study design. The tradeoff is taxonomic ambiguity.

In particular, the short-read 16S method could neither confidently nor reliably differentiate several clinically and ecologically important taxa at the species level. This is especially relevant for *Lactobacillus*, arguably the most important bacterial genus in the female urogenital microbiome, because species within this genus may differ in ecological role and clinical significance. It is also true for other taxa, including *Gardnerella* and members of the order Enterobacterales (*e.g.*, *E. coli, K. pneumoniae,* and *Proteus mirabilis*). Thus, where species level composition is required to answer a central research question, short-read 16S rRNA gene sequencing will not provide adequate resolution.

Long-read 16S rRNA gene sequencing may be particularly useful when the research question requires more precise taxonomic classification but does not necessarily justify full shotgun metagenomics. In the present study, the long-read 16S method improved taxonomic confidence and species-level assignment relative to the short-read method; for example, for members of the order Enterobacterales and of the genera *Lactobacillus* and *Gardnerella*. This makes it attractive for studies concerned with microbe-level rather than only community-level differences. However, like all amplicon-based methods, both short-read and long-read 16S rRNA gene sequencing are inherently constrained by the genomic region targeted. If closely related species differ at sequence positions not adequately captured in the amplified region, species-level discrimination may remain limited even with the longer reads. Thus, the long-read 16S method improves upon the short-read method but does not eliminate the core limitation of amplicon-based methods. Long-read shotgun metagenomic sequencing is best viewed as the most information-rich of the assessed sequencing approaches, although that does not make it universally preferable. Because it is not restricted to a targeted amplicon framework, microbe identification can, in principle, leverage sequence variation across much larger portions of the genome. In this study and related work, that capability supports species-level resolution and, in some contexts, can extend to strain-level typing (Kim, Pongpanich, and Porntaveetus 2024). The method may also provide a richer biological dataset than amplicon sequencing alone, including broader community context and the potential to detect multiple microbial types in a single test, including bacteria, archaea, fungi, DNA viruses, and plasmids. The long-read metagenomic method also can provide predicted functions, such as antibiotic resistance determinants. A practical limitation is that this method is methodologically more complex; thus, its use should be justified by whether the biological question actually requires that degree of resolution and breadth.

More broadly, these results argue against using “urine microbiome detection” as if it were a single methodological category. Future studies should specify at the outset whether the priority is viability, breadth, richness, species-level identity, absolute abundance, or reproducibility across platforms. Multi-method designs may be especially valuable in translational work, where culture and sequencing can provide complementary rather than redundant information (Hilt et al. 2014; Wolfe and Brubaker 2015; Wang et al. 2026). In addition, discordant samples deserve focused attention in future work, because they may be the most revealing with respect to the strengths, blind spots, and biological assumptions built into each workflow.

#### Clinical Implications

The clinical implications of this study are important. Our findings show that current urinary diagnostic results are shaped in meaningful ways by the method used. They do not, however, establish that detecting more microbes necessarily results in better clinical care. Analytical breadth and analytical sensitivity are not interchangeable with clinical utility.

From the clinician’s current viewpoint, the ideal detection method in most cases would be one that provides results quickly, yields the most pertinent and sufficiently encompassing data, permits clinicians to tailor the most appropriate treatment, is economical, and is readily available. In present-day practice, that treatment would often mean the shortest effective antibiotic course directed at the microbes considered most relevant to diagnosis and treatment while practicing antibiotic stewardship. From a scientific viewpoint, that formulation is preferable to one focused narrowly on classical “uropathogens,” because our findings and the broader literature suggest that the clinically relevant microbial signal may not map perfectly onto conventional pathogen categories. We are clearly not yet at the point where current diagnostic tools consistently identify the most treatment-relevant microbes across all urinary contexts, but that is the direction toward which future diagnostics should aim.

In the absence of universally agreed upon alternatives, we acknowledge that SUC will continue to be used. However, clinicians should recognize its limitations as the clearest immediate implication of our study is that SUC does not capture the full range of urinary microbes. Traditionally considered the “gold standard,” relative to the other methods, SUC performed poorly. It yielded results within 24 hours, identified isolated colonies at the species level, provided absolute abundances, and detected and identified *E. coli* and certain *Corynebacterium, Staphylococcus* and *Streptococcus* species. However, it underestimated or failed to detect fastidious, slower-growing, and rare microaerophiles, capnophiles and anaerobes, including several taxa considered to be emerging uropathogens, such as *Gardnerella* species. These results align with previous reports (Brubaker et al. 2023; Price et al. 2016; Moreland et al. 2024). These flaws are not unexpected. It was designed in the 1950s to specifically distinguish true infection from contamination and was focused specifically upon detection of pyelonephritis, especially during pregnancy (Kass 1962, 1960a, 1960b). Thus, the assay was specifically targeted towards the non-fastidious, fast-growing, facultative anaerobes that were considered to be the primary cause of that infection. As such, it is extremely biased towards a very limited range of microbes, especially *E. coli* (Moreland et al. 2024; Price et al. 2017).

In contrast, EQUC improved both sample positivity and species recovery. It was capable of detecting and identifying low-abundance bacteria, fastidious microbes, polymicrobial communities, or species not reported by standard clinical laboratory workflows. EQUC may therefore represent a practical near-term improvement over SUC in selected settings where broader viable recovery is desired. For example, a streamlined version was recommended for patients with chronic symptoms whose SUC results repeatedly return as negative.

The sequencing results extend this point but also introduce interpretive complexity. Sequencing-based methods identified many more positive samples than culture and recovered a broader taxonomic landscape. Yet the clinical meaning of that additional information is not self-evident. Detection of microbial DNA does not by itself distinguish pathogenic involvement from background community membership, transient exposure, contamination, or other biologically real but clinically neutral states. Thus, while our data indicate that current diagnostic methods differ substantially in analytical scope, they do not support the conclusion that the most analytically sensitive method is automatically the best diagnostic method. This is particularly important in urobiome contexts where broad detection may capture microbes of uncertain immediate treatment relevance.

At the same time, species-level resolution may become increasingly important as the science matures. For example, the short-read 16S method may detect the presence of *Lactobacillus* but cannot reliably distinguish among *Lactobacillus* species, even though differences among *Lactobacillus*-predominated communities may prove central to future interpretation of urinary and broader urogenital health. In that context, a method that yields only genus-level positivity may have limited clinical usefulness. Long-read approaches may become increasingly valuable where species-level precision materially affects interpretation, particularly for taxa such as *Gardnerella* or for genera in which species differ in pathogenic potential or ecological significance.

Shotgun metagenomic sequencing may ultimately provide the most comprehensive diagnostic picture, especially if future work shows that species-level discrimination, broader microbe detection, community context, calibrated microbial burden, or resistance-marker identification improve clinical interpretation, treatment selection, or outcomes. An additional conceptual advantage is that long-read metagenomic workflows may identify multiple microbial types within a single test, including bacteria, archaea, fungi, DNA viruses, and plasmids. However, the present study does not establish that this broader analytical yield changes diagnosis or management, and that translational step remains essential before any sequencing-based approach can be judged superior for routine clinical use.

The key unresolved clinical question is therefore not simply which method detects more, but which analytical features most improve clinical decisions. Those features may include viable recovery, species-level resolution, quantitative burden, polymicrobial detection, reproducibility in routine laboratory settings, time to result, and cost. Our results show that these advantages do not align perfectly within a single method. Clinical translation will therefore require endpoint-specific validation rather than technology-generic enthusiasm. Future studies should ask which methods most meaningfully improve diagnosis, treatment selection, and patient outcomes in the intended clinical population.

### Strengths and Limitations of the Study

This study has several notable strengths. First, it uses a within-sample comparative design, with multiple detection workflows applied to parallel aliquots from the same urine specimens. That design reduces one of the major challenges in microbiome methods comparisons: separating biological variation between samples from methodological variation introduced by the assay. Because each participant effectively served as their own comparator across methods, the observed differences are more plausibly attributable to the detection workflows themselves. Second, the study compares a broader range of methods than is typical in a single analysis. By including both standard and enhanced culture, as well as short-read and long-read 16S rRNA gene sequencing plus long-read shotgun metagenomic sequencing, it spans a wide portion of the currently relevant methodological landscape. Third, the analysis is multidimensional. Positivity, species recovery, alpha diversity, beta diversity, clustering behavior, differential abundance, and taxonomic confidence each provide different information, and evaluating them together yields a more realistic picture of comparative method performance than reliance on a single summary metric would. Finally, we obtained more confident taxonomic identification by combining the Bayesian-based tool BLCA with an integrated query of two different curated databases (NCBI and Greengenes2).

Our study also has important limitations. The cohort was drawn from a single academic-center obstetric population and consisted of pregnant participants undergoing screening for asymptomatic bacteriuria, with exclusions including recurrent urinary tract infection and current antibiotic use. This improves internal consistency but limits generalizability. A second limitation is denominator mismatch across methods. Of 188 total samples, only 101 underwent culture and 139 underwent long-read shotgun metagenomic sequencing. In addition, portions of the comparative analysis relied on subsets of samples positive across methods. These constraints do not invalidate our findings, but they complicate direct cross-platform interpretation and may bias some comparisons toward higher-signal or more concordant specimens. This issue matters especially because discordant specimens may be among the most informative for understanding where methods diverge analytically and where current diagnostic workflows may fail. A related concern is potential selection bias in the cultured subset and especially in analyses restricted to samples with positive signals across methods. Because SUC has been criticized for failing to grow multiple clinically or ecologically relevant genera, restricting attention to specimens that were positive across methods may bias the analysis toward a subset of urogenital communities containing one or more readily culturable organisms. As a result, this study does not fully address how many SUC-negative specimens would have returned positive microbe identifications by other methods, nor does it fully characterize the biological importance of those discordant cases. That does not undermine the value of the current comparative question, but it does narrow the interpretive reach of the conclusions. This resembles the previously recorded concern that intersection-positive subset analysis may bias comparative interpretation; the same caution should carry forward here. Finally, this is an analytical comparison study rather than an outcome-based clinical validation study. It tells us that different methods produce different microbial readouts, but it does not tell us which readout is most clinically correct or most useful. Culture and sequencing do not measure precisely the same biological thing, and disagreement between them is therefore not surprising. Culture depends on growth under defined laboratory conditions, whereas sequencing depends on nucleic acid recovery, amplification or library-preparation behavior, reference-database completeness, and computational classification. Each carries its own biases, and each may capture true but different aspects of the specimen. That should temper overly simple interpretations of concordance, discordance, or apparent superiority.

## Conclusion

In conclusion, our findings show that the urinary microbial profile obtained from a specimen depends strongly on the detection method used. EQUC improved upon SUC in positivity and species recovery, whereas sequencing-based methods broadened detection and, in some cases, improved taxonomic resolution. No single method was uniformly best across all use cases. Instead, each method contributed different strengths and different blind spots, underscoring the importance of aligning method selection with the specific research or clinical question.

For research, this means choosing methods according to whether the priority is viable recovery, broad community profiling, species-level resolution, or deeper multi-kingdom characterization. For clinical translation, it means resisting the assumption that broader detection alone is sufficient. The field now needs studies that link method-specific analytical advantages to diagnosis, treatment selection, antibiotic stewardship, and patient outcomes. Until then, the most defensible conclusion is that urinary microbial detection is method-dependent and that future progress will depend on matching the method to the purpose.

## SUPPLEMENTAL TABLES

**Supplemental Table 1.** Intersection of participants_prevalence_summary-Genus

**Supplemental Table 2.** Intersection of participants_prevalence_summary-Species

## AUTHOR STATEMENTS

### AUTHOR CONTRIBUTIONS

Conceptualization: AJW, TTP Data Curation: MK

Formal Analysis: MK, MFN

Funding Acquisition: AJW, NJP

Web Lab: BF, MJR Investigation: All authors

Project administration: AJW, NJP

Resources: AJW, NJP

Supervision: AJW, NJP

Visualization: MK

Writing – original draft: AJW

Writing – review & editing: All authors

## ETHICAL APPROVAL

This study was in ethical compliance with Loyola University’s institutional review board approval (LU216104).

## CONSENT FOR PUBLICATION

Not applicable

## ACKNOWLEDGMENTS

We would like to acknowledge the Loyola clinicians who recruited participants and collected samples, the participants themselves for consenting to the study, members of the Wolfe lab who processed the samples, the SeqCenter for the 16S rRNA gene sequencing, and Mike Fischer for his financial support.

